# Transactive response DNA-binding Protein (TDP-43) regulates early HIV-1 entry and infection

**DOI:** 10.1101/2021.12.06.471424

**Authors:** Romina Cabrera-Rodríguez, Silvia Pérez-Yanes, Rafaela González-Montelongo, José M. Lorenzo-Salazar, Judith Estévez-Herrera, Jonay García-Luis, Antonio Íñigo-Campos, Luis A. Rubio-Rodríguez, Adrián Muñoz-Barrera, Rodrigo Trujillo-González, Roberto Dorta-Guerra, Concha Casado, María Pernas, Julià Blanco, Carlos Flores, Agustín Valenzuela-Fernández

## Abstract

The transactive response DNA-binding protein (TDP-43) is an important regulator of mRNA, being reported to stabilize the anti-HIV factor, histone deacetylase 6 (HDAC6). However, little is known about the role of TDP-43 in HIV infection. In this work, we seek for the TDP-43 function on regulating CD4+ T cell permissibility to HIV infection. We observed that over-expression of wt-TDP-43 in CD4+ T cells stabilized HDAC6, increasing mRNA and the protein levels of this antiviral enzyme. Under this experimental condition, HIV-1 infection was impaired, independently of the viral envelope glycoprotein (Env) complex tropism. The results obtained by using an HIV-1 Env-mediated cell-to-cell fusion model, under the same experimental conditions, suggest that the increase in TDP-43 levels negatively affects the viral Env fusion capacity. Moreover, the specific siRNA silencing of endogenous TDP-43 in target cells lead to a significant decrease in the levels of HDAC6 which consistently induces an increase in the fusogenic and infection activities of the HIV-1 Env. These observations were confirmed by using primary viral Envs from HIV+ individuals with different clinical phenotypes. An increase in the level of expression of wt-TDP-43 strongly reduced the Envs infection activity of viremic non-progressors (VNP) and rapid progressors (RP) HIV+ individuals down to the levels of the inefficient HIV-1 Envs from long-term non-progressor elite controllers (LTNP-EC) individuals. On the contrary, low levels of endogenous TDP-43, obtained after specific siRNA-TDP-43 knocking-down, significantly favors the infection activity of primary HIV-1 Envs of VNP and RP individuals, leading to an increase in the infection ability of the primary HIV-1/LTNP-EC Envs. Based on this evidence, we interpret that TDP-43 conditions cell permissibility to HIV infection by affecting viral Env fusion and infection capacities, at least by altering the cellular levels of the antiviral enzyme HDAC6.

## Introduction

Transactive response DNA-binding protein (TDP-43) is a nuclear RNA-binding protein able to process RNA, thereby acting in different biological processes, including splicing, transcription and translation. Since TDP-43 recognizes UG-rich sequences present within approximately one third of all transcribed genes^1–3^, it is uniquely able to influence the processing of hundreds to thousands of transcripts. One of the transcripts that is mainly regulated by TDP-43 is the corresponding to the cytoplasmic enzyme, histone deacetylase 6 (HDAC6)^4^. TDP-43 bound specifically and functionally to HDAC6 mRNA. A TDP-43 knockout of *Drosophila melanogaster* has validated its biological relevance, further confirmed by the specific downregulation of HDAC6^4^. Hence, loss of functional TDP-43 causes HDAC6 downregulation and might thereby contribute to several pathogenic biological events where these two factors could be functionally involved^5–12^. Interestingly, TDP-43 contains two RNA-recognition motifs (RRMs) and a C-terminal glycine-rich domain (GRD) which was originally reported as a novel protein binding to transactivator responsive DNA sequences within human immunodeficiency virus type 1 (HIV-1) acting as a strong transcriptional repressor^13^. However, it has been reported that TDP-43 lacks any effect on LTR transcription. Thus, it did not affect HIV-1 gene expression and virus production in T cells and macrophages^14^.

It is important to note that HDAC6 is key for regulating HIV-1 infection. HDAC6 represents a barrier for viral infection and production, affecting pore fusion formation by avoiding proinfection HIV-mediated *α*-tubulin acetylation postransductional modification, and promoting HDAC6/p62 autophagy to clear key viral factors, such as Vif and Pr55Gag thereby stabilizing the HDAC6/A3G restriction factor complex^15–19^. Furthermore, there is a clear-cut correlation between the inability of primary HIV-1 envelope complex (Env) of virus from elite controller individuals (ECs) to signal and overcome the HDAC6 barrier, and this HIV+ long-term non-progressor phenotype^20^. On the contrary, functional primary Envs from virus of viremic HIV+ patients (Progressors, Rapid Progressors (RPs) and Viremic non-progressors (VNPs)) are able to overcome HDAC6 barrier^20, 21^. Therefore, it is plausible that a cellular factor that can balance cellular HDAC6 levels could regulate its anti-HIV functions and cell permissibility to HIV infection.

Altogether there is a lack of evidence involving TDP-43 in the regulation of HIV-1 viral entry an infection. We propose that due to the importance of HDAC6 for limiting HIV-1 fusion and infection in the first step of the viral cycle, TDP-43 will regulate cell permissibility to fuse and be infected by HIV-1 by acting on the levels of expression of HDAC6. Our results indicate that the TDP-43/HDAC6 axis could be crucial for regulating HIV infection.

## Materials and Methods

### Antibodies and Reagents

Rabbit anti-HDAC6 (H-300; sc-11420), and Polybrene (sc-134220) were obtained from Santa Cruz Biotechnology (Santa Cruz, CA, USA). The neutralizing mAb RPA-T4 eBioscience, San Diego, CA) directed against CD4 was phycoerythrin (PE)-labelled (for flow cytometry). PE conjugates of the CD184 (clone 12G5) and CD195 (clone 2D7/CCR5) (BD Bioscience/BD PharMingen, San Jose, CA) are directed against the second extracellular loop of CXCR4 and CCR5, respectively. Secondary Alexa Fluor 568-goat anti-mouse (to label TDP-43) and Alexa Fluor 488-goat anti-rabbit (to label HDAC6) were purchased from Molecular Probes (Eugene, OR). Rabbit anti-TDP-43 (Catalog Number T1705); Mouse anti-TDP-43 mAb (clone 2E2-D3, catalog Number WH0023435M1); Mouse Monoclonal anti-Flag M2 (F1804); Mabs anti-α-tubulin (T6074) and anti-acetylated *α*-tubulin (T7451); and secondary horseradish peroxidase (HRP)-conjugated Abs, specific for any Ab species assayed were purchased from Sigma-Aldrich (Sigma-Aldrich, St. Louis, USA). Complete™ Protease Inhibitor Cocktail (11697498001) was obtained from Roche Diagnostics (GmbH, Mannheim, Germany).

### DNA plasmids and viral DNA constructs

Vectors for expression of wild-type TDP-43 (Flag-wt-TDP-43) or mutant at the import NLS signal (Flag-NLS-mut-TDP-43) were courtesy of Dr. Thorsten Schmidt^4^. The pNL4-3.Luc.R-E-provirus (*Δnef*/*Δenv*), the R5-tropic BaL.01-*envelope* (*env*) glycoprotein plasmid (catalog numbers 6070013 and 11445, respectively), the X4-tropic HXB2-*envelope* (*env*, catalogue number 5040154), and the pHEF-VSV-G vector (catalogue number 4693) encoding the vesicular stomatitis virus G (VSV-G) protein were obtained via the NIH AIDS Research and Reference Reagent Programme.

### Samples and *env* clones and ethics statement

Samples from virus of HIV+ RP, LTNP-EC and VNP individuals were obtained and studied as previously described^20, 21^. Samples were obtained from participants in previous studies who gave informed consent for the study. Consents were approved by the Ethical and Investigation Committees for their corresponding hospital and the samples were encoded and de-identified in these centers. All clinical investigations were conducted according to the principles expressed in the Declaration of Helsinki in 1975, as revised in 1983. For LTNP-EC Envs the studies were approved by the “Comité de ética de la Investigación y de Bienestar Animal of the Instituto de Salud Carlos III” under no. CEI PI 05_2010-v3 and CEI PI 09-2013v2, and for VNP and RP Envs we confirm that all methods were carried out in accordance with relevant guidelines and regulations. All procedures were approved by the Ethics committee of the Hospital Germans Trias I Pujol. All individuals provided their written informed consent.

### Cells

The human CEM.NKR-CCR5 permissive cell line (catalog number 4376, NIH AIDS Research and Reference Reagent Program) and the HEK-293T cells (catalog number 103, NIH AIDS Research and Reference Reagent Program) were grown at 37°C in a humidified atmosphere with 5% CO2 in RPMI 1640 medium (Lonza, Verviers, Belgium) in the case of the CEM.NKR-CCR5 cells, and in DMEM (Lonza) when HEK-293T cells were used, both medium supplemented with 10% fetal calf serum (Lonza), 1% L-glutamine, and 1% penicillin-streptomycin antibiotics (Lonza) and mycoplasma free (Mycozap antibiotics, Lonza), and both cell lines were regularly splitted every 2-3 days. The HEK-293T cell line was cultured to 50-70% confluence in fresh supplemented medium 24 h before cell transfection with viral or human DNA constructs. The HeLa-P5 cells are constitutively expressing CXCR4 and stably transfected with human CD4 and CCR5 cDNAs and with an HIV long terminal repeat-driven-β-galactosidase reporter gene^15, 22, 23^, as well as, HeLa-243 and HeLa-ADA cells, co-expressing the Tat and X4- and R5-tropic HIV-1-Env proteins, respectively, were provided by Dr. M. Alizon (Hôpital Cochin, Paris, France) and cultured as previously described^15, 22, 23^. The HEK-293T cell line was similarly cultured in supplemented DMEM (Lonza), as reported^16, 17, 20, 21, 24^. Cells were harvested and cultured to 50-70% confluence in fresh supplemented DMEM 24 h before cell transfection with viral or human DNA constructs.

### Western Blotting

Protein expression was determined by SDS-PAGE and Western blotting in cell lysates. Plasmids or short interference RNA (siRNA) oligonucleotides (oligos) were nucleofected into CEM.NKR-CCR5 cells with Amaxa kits C (1 μg into 2×10^6^ cells/mL) or transfected into HeLa P5 cells with polyethylenimine (PEI). PEI with an average molecular mass of 25 kDa (PEI25K) (Polyscience, Warrington, PA, United States) was gently vortexed, incubated for 20-30 min at room temperature (RT), and then added to cells in culture. Briefly, 24 h after nucleofection in the case of overexpression of constructs and 48 h after nucleofection in the case of siRNA oligos, cells were lysated in lysis buffer (1% Triton-X100, 50 mM Tris-HCl pH 7,5, 150 mM NaCl, 0.5% sodium deoxycholate, and protease inhibitor (Roche Diagnostics), for 30 min and sonicated for 30 s at 4°C. Equivalent amounts of protein (30-40 µg), determined using the bicinchoninic acid (BCA) method (Millipore Corporation, Billerica, MA), were resuspended and treated by Laemmli buffer and then were separated in 10% SDS-PAGE and electroblotted onto 0.45 µm polyvinylidene difluoride membranes (PVDF; Millipore) using Trans-blot Turbo (Bio-Rad, Hercules, CA). Membranes were blocked 5% non-fat dry milk in TBST (100 mM Tris, 0.9% NaCl, pH 7.5, 0.1% Tween 200) for 30 min and then incubated with specific antibodies. Proteins were detected by luminescence using the ECL System (Bio-Rad), and analyzed using a ChemiDoc MP device and Image LabTM Software, Version 5.2 (Bio-Rad).

### Immunofluorescence and confocal microscopy analysis

Control or transfected Lymphocytes (with Flag-wt-TDP-43 or Flag-NLS-mut-TDP-43 plasmid (1 μg) using nucleofection kit (Lonza)) were placed on coverslips (2 x 10^6^ cells in sterile glass coverslips-Ø 12 mm) to be immunolabeled and analyzed by confocal microscopy 48 h post-transfection. Then, cells were washed three times with PBS and fixed for 20 min in 2% paraformaldehyde with 1% sucrose in PBS. Then, cells were washed three times with PBS and permeabilized with 0.1% Triton X-100 in PBS. These cells were then washed with PBS and immunostained for 1 h at room temperature (RT) by Alexa 568-labeled goat anti-mouse against Flag-TDP-43 constructs (wt and NLS-mut) previously incubated with a specific mouse mAb (anti-TDP-43 or anti-Flag) and diluted in PBS with 0.1% BSA. Secondary Alexa Fluor 568-goat anti-mouse was used to label TDP-43 (endogenous or Flag-constructs previously bound to mouse anti-TDP-43 or mouse anti-Flag M2 mAb), and Alexa Fluor 488-goat anti-rabbit was used to label endogenous HDAC6 (previously bound to rabbit anti-HDAC6 Ab). Dapi was used to stain the nucleus of cells. Then, coverslips carrying these cells were mounted in Mowiol-antifade (Dako, Glostrup, Denmark) and imaged in x-y mid-sections in a FluoView FV1000 confocal microscope through a 1.35 numerical aperture (NA) objective (60×) (Olympus, Center Valley, PA) for high-resolution imaging of fixed cells. The final images and molecule codistributions were analyzed with MetaMorph software (Universal Imaging, Downington, PA) as previously reported^24–28^.

### Messenger RNA silencing

We used the below indicated short interference RNA oligonucleotides (siRNA), specifically directed against the indicated mRNA sequences of TDP-43, in order to knockdown TDP-43 expression. Transient siRNA transfections were performed using Amaxa kits C (Amaxa Lonza) for CEM.NKR-CCR5 cells, or polyethylenimine when using HeLa P5, with 1 μM of a commercial scramble control or TDP-43-specific siRNAs: siRNA-TDP-43 A: 5’-CAAUAGCAAUAGACAGUUA[dT][dT]-3’; siRNA-TDP-43 B: 5’-CACUACAAUUGAUAUCAAA[dT][dT]-3’; siRNA-TDP-43 C: 5’-GAAUCAGGGUGGAUUUGGU[dT][dT]-3’; siRNA-TDP-43 D: 5’-GAAACAAUCAAGGUAGUAA[dT][dT]-3’ (Sigma)^4^. The siRNA for TDP-43 induced specific interference of protein expression for at least 72 h, as witnessed in Western blots (data not shown). Cells nucleofected with scrambled or specific siRNA oligos against TDP-43 were lysed and analyzed with specific antibodies in Western blots to determine the endogenous TDP-43 silencing.

### RNA extraction and RT-qPCR

RNA from CEM.NKR-CCR5 cells was isolated using the RNeasy Mini kit (Qiagen) following manufacturer’s instruction. The cellular qRT–PCR, 1000 ng total RNA was reverse transcribed using iScript™ cDNA Synthesis Kit (Bio-Rad), random primers and anchored oligo-dT primer. RT reaction (20 μL) was used as template for transcript amplification; 1/10 dilutions were used in triplicates with 0.2 μM primer and 10 μL LightCycler 480 SYBR Green I Master in a 20 μL reaction and qPCR executed in a 98- well block on a CFX96 Touch Real-Time PCR Detection System (Bio-Rad). Absolute transcript levels for TDP-43, HDAC6 and the controls were obtained by second derivative method. Relative transcript levels were calculated as TDP-43 or HDAC6/controls ratio and normalized to the relative expression level of the mock-transfected control. The primers used for qPCR were as follows: for HDAC6 (Forward: 5′- ATGCAGCTTGCGGTTTTTGC -3’ / Reverse: 5′- TGCTGAGTTCCATTACCGTGG -3’); for TDP-43 (Forward: 5’- GCTTCGCTACAGGAATCCAG -3’/ Reverse: 5’- GATCTTTCTTGACCTGCACC - 3’); and for GADPH as housekeeping gene (Forward: 5’- GGAAGCTCACTGGCATGGCCT -3’ / Reverse: 5’- CGCCTGCTTCACCACCTTCTTG -3’).

### RNA sequencing

#### Library preparation and sequencing

Libraries were prepared using 50 ng of total RNA (all with RIN > 8.5) according to manufacturer instructions with the Illumina® Stranded Total RNA Prep, Ligation with Ribo-Zero Plus (Illumina, Inc.), including the depletion of cytoplasmic, mitochondrial, and beta globin rRNA. After depletion, libraries were prepared with 50 ng of total RNA. Library integrity and size (size range of 360 to 400 bp) was confirmed by analysis on a 4200 TapeStation (Agilent) with D1000 ScreenTape. Library concentrations were determined using the dsDNA HS assay kit on Qubit 3.0 (Thermo Fisher Scientific). Sequencing was performed with paired-end 75 cycles on a NextSeq 550 System (Illumina, Inc.) at the Instituto Tecnológico y de Energías Renovables.

#### Bioinformatic analysis

The raw sequencing data was demultiplexed with bcl2fastq v2.20 and an initial assessment was performed with FastQC v0.11.8. Salmon v1.5.0^29^ was used to quantify the abundance (TPM, transcripts per million) of expression at the transcript level, using the GRCh37/hg19 as the reference. In order to make an assessment on the mapping quality, STAR v2.7.9a^30^ was used to align reads to reference, Qualimap v2.2.1^31^ was used to compute quality metrics, and MultiQC v1.10.1^32^ to aggregate the results, QC metrics, and sample data. Finally, differential gene expression analysis was computed with DESeq2 v.1.32.0^33^ package for R. All computations were performed in the TeideHPC supercomputing infrastructure (http://teidehpc.iter.es/en).

### Cell fractioning

For the cell fractioning, two buffer extraction were used, one as Cytosolic Extraction Buffer (CEB) (HEPES 10 mM; KCl 60 mM; EDTA 1 mM; NP40 1%) and another one as Nuclear Extraction Buffer (NEB) (HEPES 20 mM; KCl 500 mM; MgCl2 1.5 mM; EDTA 0.2 mM, glycerol 25% v/v). First, 2×10^6^ CEM.NKR-CCR5 cells were washed with 100 μL PBS and centrifuge at 1500 rpm (200 x *g*) x 5 min. Then, the pellet was resuspended (by gentle pipetting) in CEB, usually about 30-50 μL (5X times the volume of pellet) and put into 4°C for 20 min. Centrifugation was done again at 10000 rpm (12000 x *g*) for 10 min at 4°C, and the supernatant was collected as the cytosolic fraction. The pellet (nuclear fraction) was washed at least twice with CEB to eliminate cytoplasmic debris (centrifugation between washes could be for 5 min 4°C at 10000 rpm (12000 x *g*)), then resuspended in 60 μL NEB (having in account the nucleus-cytoplasm ratio) and incubated at 4°C for 30 min, vortexing once every 10 min. At last, the pellet was centrifuged 15 min, 4°C at 12,000 x *g*, and collected supernatant as the nuclear fraction. For Western Blot analysis, the *α*-tubulin was used as the cytoplasmic marker, and Lamin B1 as the marker for the nuclear fraction.

### HIV-1 Env mediated cell-to-cell fusion assay

A β-Galactosidase cell fusion assay was performed as previously described^24, 28^. Briefly, HeLa-243 or HeLa-ADA cells were mixed with HeLa-P5 cells or control, or previously transfected with TDP-43 constructs or siRNA-TDP-43 oligos, in 96-well plates in a 1:1 ratio (20,000 total cells). These cocultures were kept at fusion for 16 h at 37°C. The fused cells were washed with Hanks’ balanced salt solution and lysed, and the enzymatic activity was evaluated by chemiluminescence (β-Gal reporter gene assay, Roche Diagnostics). Anti-CD4 neutralizing mAb L3T4 was used as a control for the blockage of cell fusion (5 µg/mL was preincubated in HeLa-P5 cells for 30 min at 37°C before co-culture with Env+-HeLa cells).

### Production of viral particles with luciferase-reporter pseudoviruses

Replication-deficient luciferase-HIV-1 viral particles (luciferase-reporter pseudoviruses) were obtained as described^15–17, 20, 21, 24, 28^, using the luciferase-expressing reporter virus HIV/*Δnef*/*Δenv*/*luc*+ (pNL4-3.Luc.R-E-provirus bearing the *luciferase* gene inserted into the *nef* ORF and not expressing *env*; catalog number 6070013, NIH AIDS Research and Reference Reagent Program), and the described *env* expression plasmids from the different HIV-1 individuals or the reference *env* plasmids studied. Thus, HIV-1 viral particles were produced in 12-wells plates by co-transfecting HEK-293T packaging cells (70% confluence) with pNL4-3.Luc.R-E- (1 µg) and R5-tropic (BaL.01), or the X4-tropic HXB2 or a primary Env-glycoprotein vector (1 µg). Viral plasmids were transduced in HEK-293T cells using X-tremeGENE HP DNA transfection reagent (Roche). After the addition of X-tremeGENE HP to the viral plasmids the solution was mixed in 100 µL of DMEM medium without serum or antibiotics, and incubated for 20 min at RT prior to adding it to HEK-293T cells. The cells were cultured for 48 h to allow viral production. After this, viral particles were harvested. Viral stocks were normalized by p24-Gag content as measured with an Enzyme Linked Immunosorbent Assay test (GenscreenTM HIV-1 Ag Assay; Bio-Rad, Marnes-la-Coquette, France). Virions were used to infect CEM.NKR-CCR5 cells after ELISA-p24 quantification and normalization. As a control for efficiency of viral production, a CD4 independent viral entry and infection assay was performed in parallel by co-transducing the pNL4-3.Luc.R-E-vector (1 μg) with the pHEF-VSV-G vector (1 μg; National Institutes of Health-AIDS Reagent Program), thereby generating non-replicative viral particles that fuse with cells in a VSV-G-dependent and CD4- independent manner^34^.

### Luciferase viral entry and infection assay

CEM.NKR-CCR5 cells (9×10^5^ cells in 24-well plates with 20 μg/mL of Polybrene) were infected with 200 ng of p24 of luciferase-reporter pseudoviruses, bearing R5- tropic BaL (BaL.01-*env* plasmid, catalog number 11445, NIH AIDS Research and Reference Reagent Program), X4-tropic HXB2 (*env*, catalogue number 5040154, NIH AIDS Research and Reference Reagent Program) or primary Envs, in 1 mL total volume with RPMI 1640 for 2 h (by centrifugation at 1,200 g at 25°C) and subsequent incubation for 4 h at 37°C, as previously described^15–17, 20, 21, 24, 25, 28, 34^. Unbound viruses were then removed by washing the infected cells. After 24 h of infection, luciferase activity was measured using a luciferase assay kit (Biotium, Hayward, CA) with a microplate reader (VictorTM X5; PerkinElmer, Waltham, MA).

### Flow cytometry analysis

Overexpression effects of the constructs Flag-wt-TDP-43 or Flag-NLS-mut-TDP-43 or the siRNA-TDP-43 on CD4, CCR5 and CXCR4 cell-surface expression in permissive CEM.NKR-CCR5 cells or transfected into HeLa P5 cells, were studied by flow cytometry analysis. Briefly, 24 h nucleofected/transfected cells with fluorescent constructs were incubated in ice-cold (+4°C) PBS buffer, with an anti- CD4/CCR5/CXCR4 antibody coupled to PE. Labelling of cell-surface receptors was performed by staining with a PE-conjugated IgG isotype. Cells were then washed by ice cold PBS, fixed in PBS with 1% paraformaldehyde and analyzed by flow cytometry (BD Accuri™ C6 Plus Flow Cytometer, BD Biosciences), measuring cell-surface CD4, CCR5 and CXCR4 receptors labelling as described elsewhere^15, 17, 24, 25, 28, 34^. Basal cell fluorescence intensity for receptors labelling was determined by staining cells with a PE-conjugated IgG isotype control in cells over expressing free pCDNA3.1(-) empty vector.

### Statistical analysis

Statistical analyses were performed using GraphPad Prism, version 6.0b (GraphPad Software). Significance when comparing groups was determined with a 2-tailed nonparametric Mann-Whitney U-test.

## Results

### Characterization of the expression and cellular localization of full-length and a nuclear localization signal mutant of TDP-43

To study the role of TDP-43 in the control of HIV-1 infection, we first analyzed the pattern of expression of TDP-43 in permissive CEM.NKR-CCR5 T cells. We observed that transiently over-expressed Flag-wt-TDP-43 (N-terminal Flag-tagged, wild-type (wt) TDP-43 construct) mainly localizes in the isolated nuclear fraction of cells as observed by biochemical detection (**Figure 1**, *Flag associated western-blot bands*), being also detected in the cytoplasmic fraction, similarly as observed for the endogenous TDP-43 (**Figure 1**, *lower molecular weight TDP-43 western-blot bands*). As a control, we used a N-terminal Flag-tagged NLS-TDP-43 mutant (NLS-mut-TDP-43) that lacks the nuclear localization signal (NLS) of the protein^4, 35, 36^. This mutant is mainly located at the cytoplasm, as observed by biochemical detection of cell fractioning (**Figure 1A**, *Flag and TDP-43 associated western-blot bands*). The quantification of the level of expression of these two TDP-43 constructs is shown in **Figure 1B**. Similar results are obtained when these cells are analyzed by fluorescent confocal microscopy (**Figure 1C**). Endogenous TDP-43 and Flag-wt-TDP-43 are both distributed in the nucleus (**Figure 1C**, *see associated series of images*), whereas the TDP-43 mutant is mainly located at the cytoplasm (**Figure 1C**, *Flag-NLS-mut-TDP-43 series of images*).

**Figure 1.**
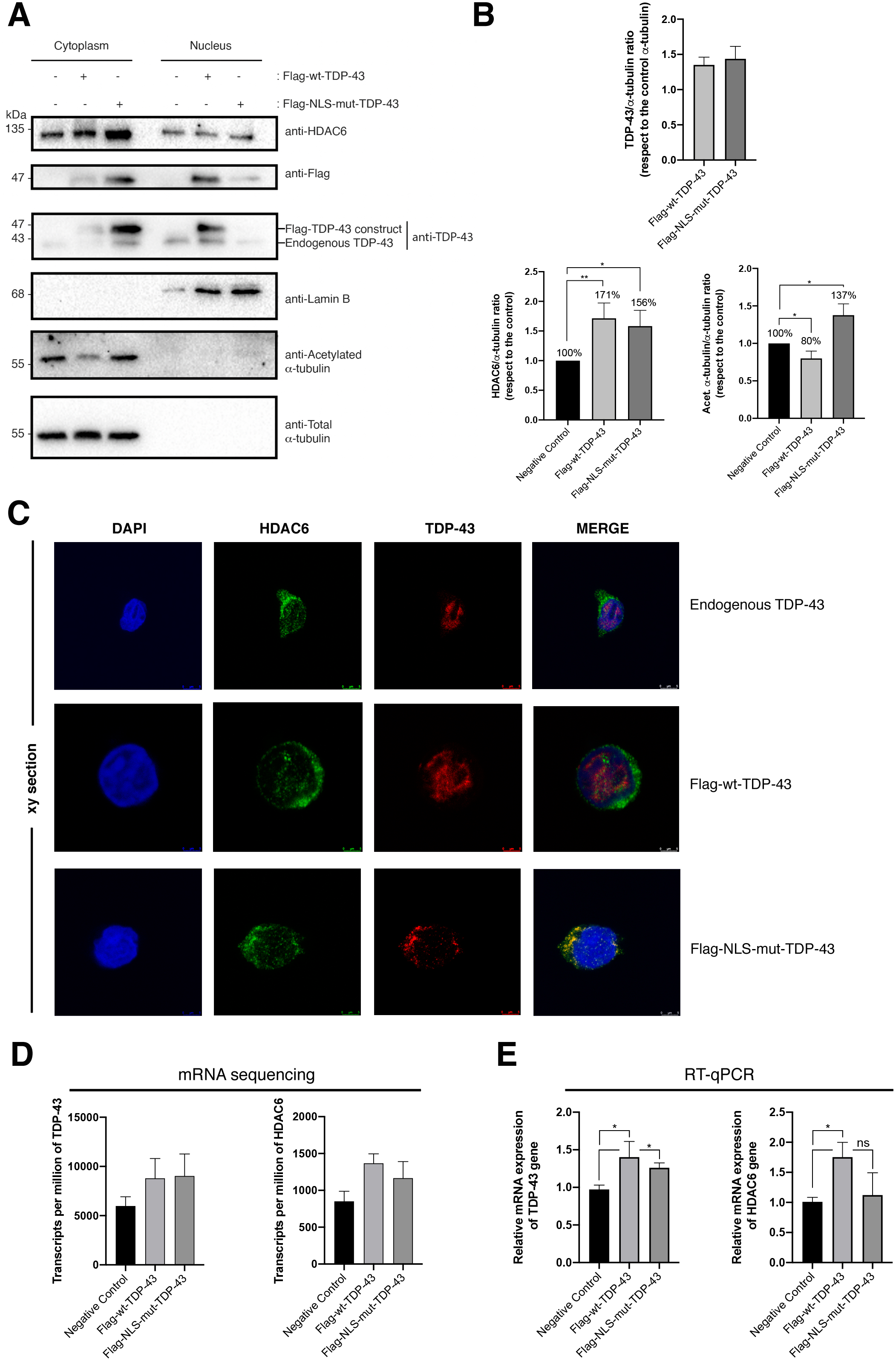
Overexpression of TDP-43 constructs regulates HDAC6 mRNA and protein stability and favors its deacetylation function. **(A** and **B)** Quantitative western blot analysis of cell fractioning after overexpression of Flag-wt-TDP-43 or Flag-NLS-mut-TDP-43 constructs (1 μg cDNA each) (**A**) and their effect over endogenous HDAC6 and *α*-tubulin acetylation. HDAC6/*α*-tubulin, acetylated *α*-tubulin/*α*-tubulin and Flag/*α*-tubulin intensity band ratios are shown (**B**), where high acetylated *α*-tubulin levels are a read-out for MTs stabilization after HDAC6-deacetylase activity decreasing. A representative experiment of six performed in CEM.NK-CCR5+/CD4+ T cells is shown. Histograms show intensity bands quantitation of top western blots for Flag-wt-TDP-43 or Flag-NLS-mut-TDP-43, normalized by the total amount α-tubulin, under any experimental condition. Data are means ± standard errors of the means (S.E.M.) of six independent experiments. Lamin B1 in the protein control for the nuclear fraction and *α*-tubulin for the cytoplasm fraction. When indicated, *P < 0.01 and **P < 0.05 are P values for Student’s *t* test. a.u., arbitrary light units. **(C)** Fluorescence confocal microscopy analysis of CEM.NK-CCR5+/CD4+ T cells nucleofected with both TDP-43 constructs, in order to analyse the distribution of the nucleofected proteins. 48 h post-nucleofection, fixed cells were imaged in **xy** mid-sections in a confocal microscopy (Leica TCS SP5; Leica Microsystems, Wetzlar, Germany) in a 1.35 NA objective (60x). Endogenous TDP-43 is located at the nucleus of the cell. Flag-wt-TDP-43 is mainly distributed to the nucleus with some cytoplasmic distribution. However, the Flag-NLS-mut-TDP-43 construct is excluded form the nucleus, presenting cytoplasmic localization. Endogenous HDAC6 is mainly located at the cytoplasm and presenting nuclear distribution too. The nucleus is displayed by DAPI labelling. A series of merge images are shown for any experimental condition. **(D** and **E)** Relative mRNA quantification in transcripts per million by sequencing (by duplicate) (**D**) or by RT-qPCR (by quintuplicate) (**E**) of TDP-43 and HDAC6 genes is represented in histograms under overexpression of TDP-43 constructs. When indicated, *P < 0.01 and **P < 0.05 are P values for Student’s *t* test.

### Full-length TDP-43 stabilizes HDAC6 and diminishes α-tubulin acetylation

We observed that in these HIV permissive CEM.NKR-CCR5 T cells the over-expression of TDP-43 constructs enhance the protein level of HDAC6 (**Figures 1A** and **1B,** *western-blot bands* and *associated histograms*, respectively). It is thought that TDP-43 stabilizes HDAC6 cells levels by acting on mature mRNA of HDAC6^4^. In this matter, we have observed the stabilization of higher levels of mRNA of HDAC6 in cells over-expressing the TDP-43 constructs compared to control cells, by quantification of isolated and specifically sequenced mRNA of HDAC6 (**Figures 1D** and **1E**). By this technique we determined that we have over-expressed similar levels of the two TDP-43 constructs (**Figures 1D** and **1E**), as confirmed by biochemical detection of the associated proteins (**Figures 1A** and **1B**).

The increase in the level of the HDAC6 tubulin-deacetylase enzyme, by over-expression of wt-TDP-43, induces a diminution in the level of acetylation of *α*-tubulin subunit of the microtubules (MTs) (**Figure 1A**, *acetylated a-tubulin in the cytoplasmic fraction*), a post-translational modification associated to stable MTs^37–44^. This effect is not observed by the over-expression of the NLS-mut-TDP-43 construct (**Figure 1A**). Perhaps its cytoplasmic localization may alter the HDAC6 tubulin-deacetylase activity, as it has been reported by the interaction of TDP-43 with p62 and HDAC6 in the cytoplasm^45–50^. Here, we have used this mutant construct as a control for studying TDP-43 cellular localization. As it is reported, we observed that over-expression of the wt-TDP-43 construct stabilizes mRNA and protein levels of the HDAC6 enzyme, which in turn promotes the deacetylation of acetylated *α*-tubulin as we describe here. Therefore, we seek for the TDP-43 function on regulating cell permissibility to HIV infection by working with wt-TDP-43 construct or by specific siRNA-TDP-43 silencing.

### TDP-43 overexpression inhibits HIV-1 entry and infection

First, we studied the effect of wt-TDP-43 on the early infection of permissive CEM.NKR-CCR5 T cells with HIV-1. As shown in **Figure 1**, cells over-expressing wt-TDP-43 were infected by using HIV-1, luciferase-reporter pseudovirus bearing a reference CCR5 tropic (R5-tropic) BaL envelope (Env) complex or an X4-tropic HXB2 Env (a reference CXCR4 tropic Env). We observed that over-expression of wt-TDP-43 did not significantly changes the expression of CD4, CCR5 and CXCR4 (i.e., main receptor and co-receptors for HIV infection, respectively) (**Figure 2A**). The cell infection by using pseudovirus bearing BaL or HXB2 Env was importantly diminished in cells over-expressing wt-TDP-43 compared to control, untreated cells (**Figure 2B**). Therefore, the reduction on the permissibility to HIV-1 infection of this CD4+ T cells appears to be independent of the viral tropism.

**Figure 2.**
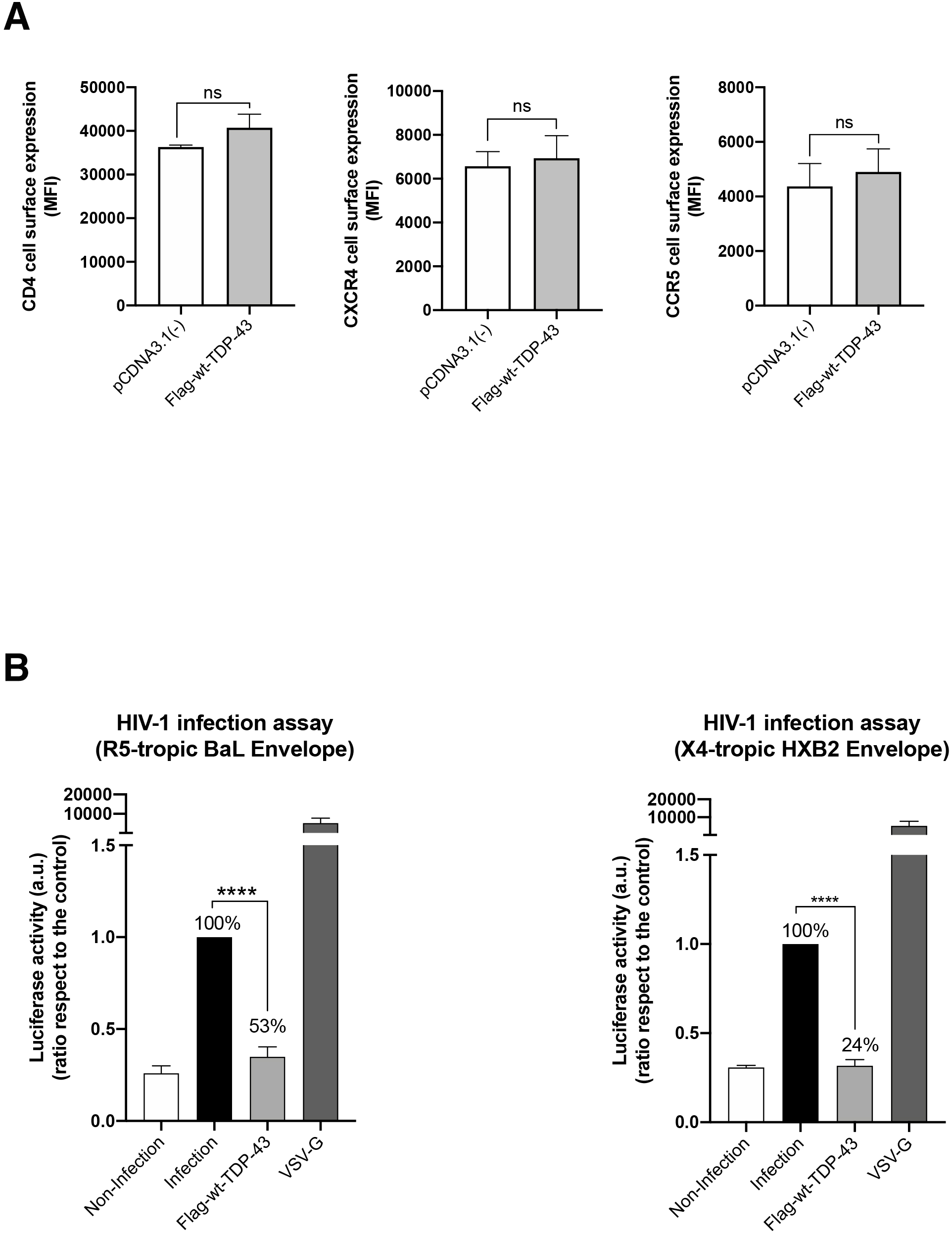
Analysis of CD4 receptor, CXCR4 and CCR5 co-receptors on the cell membrane surface and virus infection with reference envelopes after overexpression of Flag-wt-TDP-43 or Flag-NLS-mut-TDP-43 constructs. **(A)** Effect of overexpression of Flag-wt-TDP-43 or Flag-NLS-mut-TDP-43 constructs in CEM.NK-CCR5+/CD4+ T cell according to cell-surface expression of viral receptors was analyzed under each experimental condition. Data are mean ± S.E.M. of four independent experiments carried out in triplicate * p < 0.05, ** p < 0.01, *** p < 0.001, Student’s t-test. **(B)** Effects of both TDP-43 constructs in HIV-1 infection with Env reference pseudoviruses. Viral particles produced with R5-tropic pNL4-3.Luc.R-E-/Env-BaL or /X4-tropic Env-HBX2 viral inputs with CEM.NK.CCR5 permissive T cells previously nucleofected with TDP-43 constructs, and assayed in luciferase-quantified viral infection experiments. Data are mean ± S.E.M. of six independent experiments. VSV-G pseudotyped viruses were used as control of infective viral production and infection.

The TDP-43-mediated increase in the level of the HDAC6 enzyme and the subsequent diminution of the levels of acetylated *α*-tubulin in MTs (**Figure 1**) promote a cellular environment that limits early HIV-1 infection, as previously reported^15, 20, 21^. Hence, over-expression of wt-TDP-43 negatively regulates early viral entry an infection, an antiviral activity that at least accounts to the associated increase of the HDAC6-tubulind deacetylase antiviral action.

### TDP-43 over-expression inhibits HIV-1 Env-mediated pore fusion formation

As reported, HDAC6 tubulin-deacetylase activity limits HIV-1 Env-mediated pore fusion. HDAC6 impairs HIV-1-mediated stabilization of a cortex of stable and acetylated MTs in its *α*-tubulin subunit, which is required for efficient viral entry and infection, and is directly dependent on the first viral Env/CD4 interaction and signaling^15, 20, 21^. Hence, we next studied the effect of wt-TDP-3 in the control of HIV-1 Env-mediated pore fusion formation in an Env-mediated cell-to-cell fusion model^15, 22, 23, 28, 34^ (see Materials and methods). In target HeLa P5 cells, the over-expression of wt-TDP-43 did not negatively affect the expression of CD4, CCR5 and CXCR4 at cell-surface (**Figure 3A**), and increased the level of the HDAC6 protein, subsequently diminishing the level of acetylated *α*-tubulin (**Figure 3B**). Under these experimental conditions, our results indicate that co-culture of these HeLa P5 with HeLa cells expressing 243 Env (X4-tropic) or ADA Env (R5-tropic) led to a reduction of HIV-1 Env-mediated cell-to-cell fusion, being a direct measurement of the inhibition of HIV-1 Env-mediated pore fusion formation in cells over-expressing wt-TDP-43 (**Figure 3C**).

**Figure 3.**
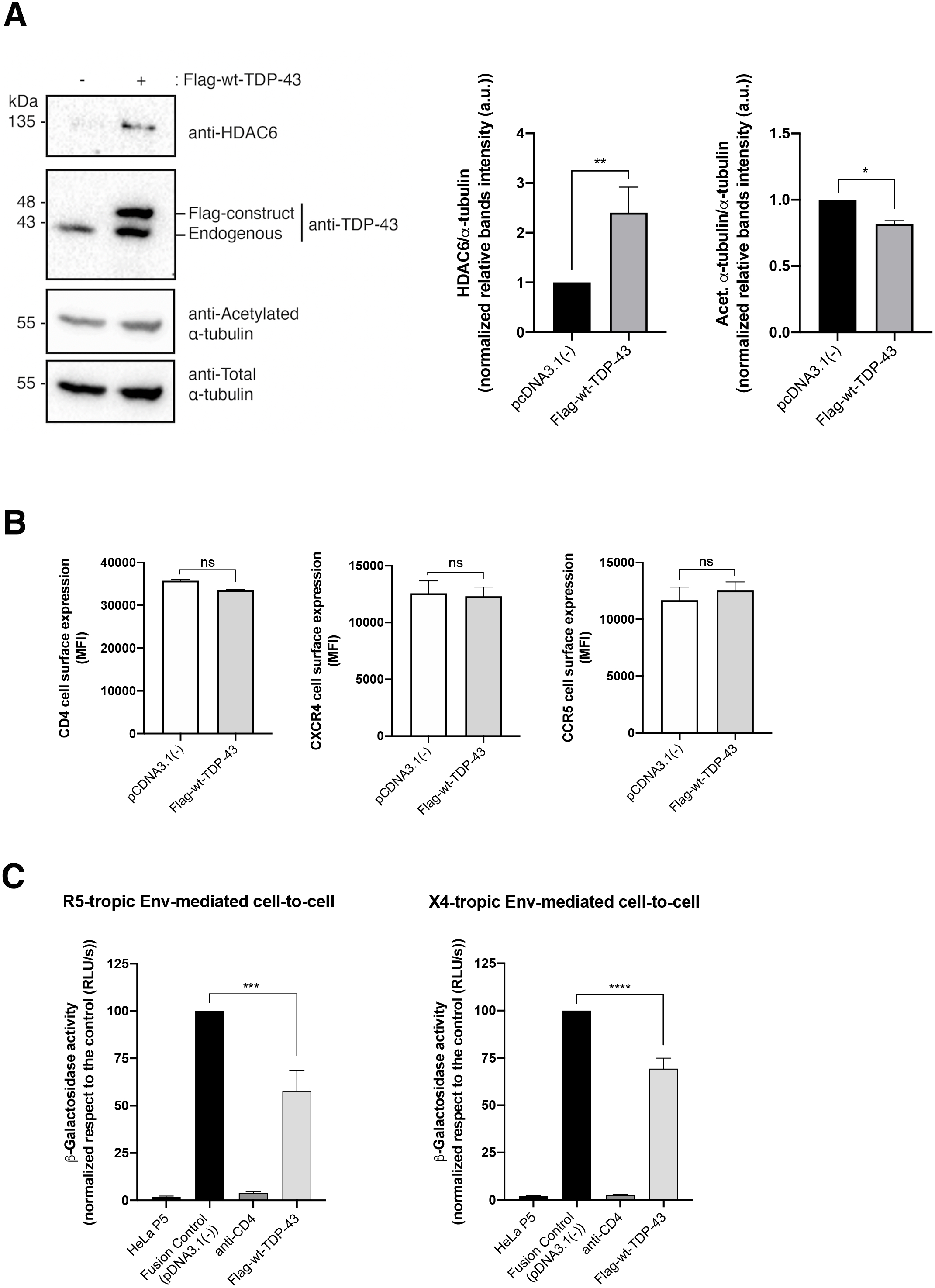
Membrane fusion is impaired in the presence of TDP-43 constructs. **(A)** Quantitative western blot analysis of HeLa. P5 cell lysate after overexpression of Flag-wt-TDP-43 construct, using 1 μg and its effect over endogenous proteins (HDAC6 and acetylated *α*-tubulin). HDAC6/*α*-tubulin and acetylated *α*-tubulin/*α*-tubulin intensity band ratios are shown. **(B)** Effect of overexpression of Flag-wt-TDP-43 construct in HeLa P5 according to cell-surface expression of viral receptors was analyzed under experimental condition. Data are mean ± S.E.M. of four independent experiments carried out in triplicate * p < 0.05, ** p < 0.01, *** p < 0.001, Student’s t-test. **(C)** Effect of Flag-wt-TDP-43 construct in HeLa P5 cells on HIV-1 Env-mediated membrane fusion measured by β-Galactosidase production, and referred to the pcDNA3.1(-) control condition (100% of membrane fusion). Untreated HeLa P5 cells were used as non-fused cells control. Treated HeLa P5 cells were fused with HeLa cells expressing X4-tropic (243) or R5-tropic (ADA) Env, in the presence or absence of a neutralizing anti-fusogenic anti-CD4 mAb.

Altogether these results indicate that the efficiency of HIV-1 infection and Env-mediated cell fusion is inversely related to HDAC6 activity which is under the control of TDP-43.

### Specific siRNA silencing of endogenous TDP-43 decreases HDAC6 protein level, increase α-tubulin acetylation and enhances cell permissibility to HIV infection

We checked four different siRNA-TDP-43 oligos (i.e., siRNA-TDP-43 A to D; **Figure 4A**) previously reported^4^, in order to assay their ability to specifically interfere the mRNA of TDP-43 and knockdown the protein. Three of them present good activity to specifically silence TDP-43 (**Figure 4A**, *siRNA-TDP-43 A, B and C oligos-associated western blot bands* and **Figure 4B**, *quantified in associated histogram*). Moreover, we quantified the amount of the mRNA of TDP-43 and HDAC6 present in cells nucleofected with the four siRNA-TDP-43 oligos and some mixture combination of them (**Figure 4C**), by direct quantification by sequencing of their mRNAs, and also by RT-qPCR (see Materials and methods) (**Figure 4C** and **4D**, respectively).

**Figure 4.**
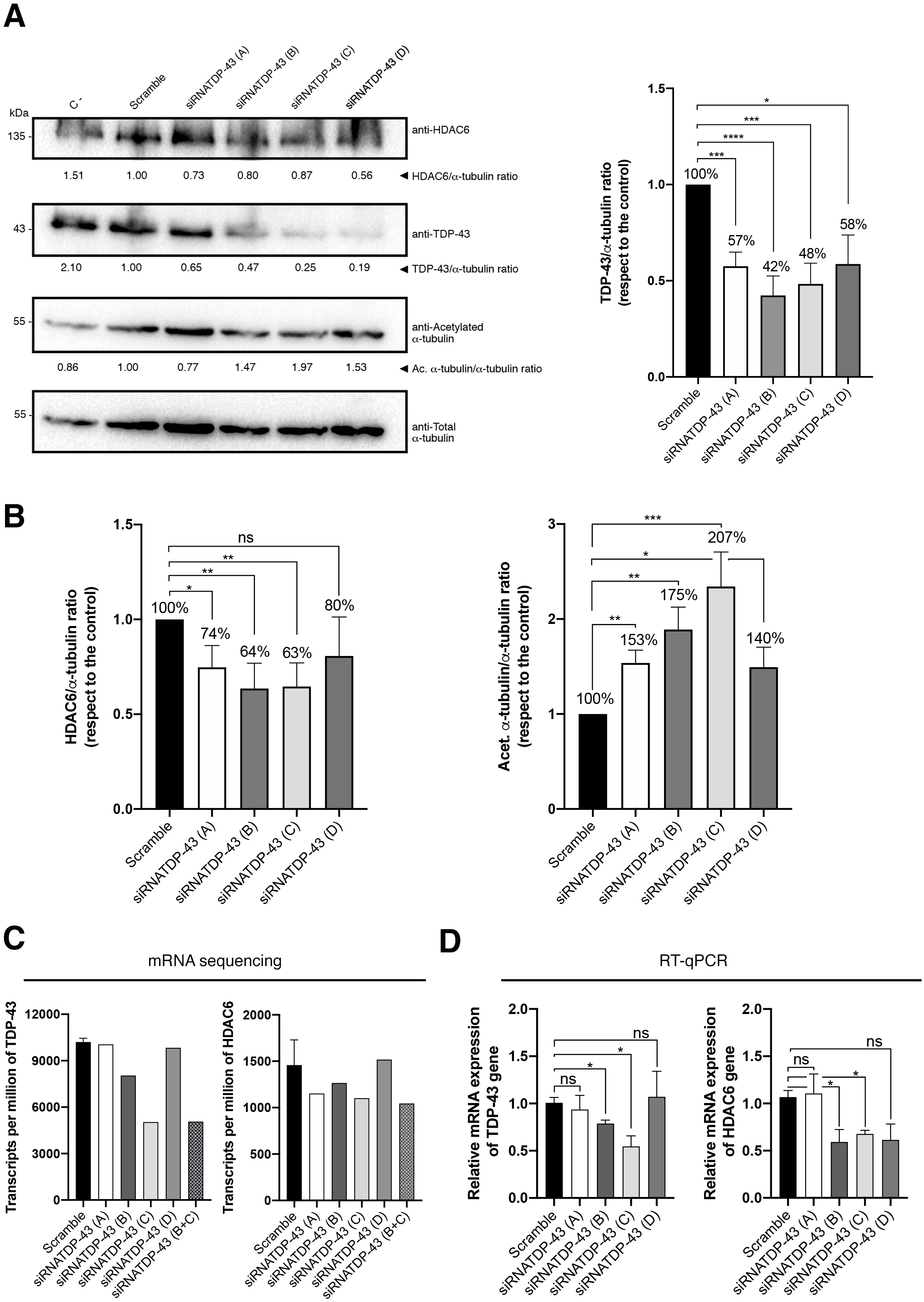
TDP-43 mRNA silencing downregulates mRNA and protein of HDAC6 impairing its deacetylation functions. **(A)** Western blot analysis of endogenous TDP-43 knockdown in permissive CEM.NK-CCR5+/CD4+ T cell nucleofected with each of the four siRNA-TDP-43 oligos used. Cells nucleofected with scrambled siRNA were used as control. One representative experiment of four is shown. Protein quantification respect to the control α-tubulin protein is shown in the histogram for TDP-43 (*right panel histogram*). **(B)** Protein quantification respect to the control α-tubulin protein is shown in the histograms for HDAC6, TDP-43 and acetylated *α*-tubulin. **(C** and **D)** Relative mRNA quantification in transcripts per million by sequencing (without replicates) (**C**) or by RT-qPCR (by quintuplicate) (**D**) of TDP-43 and HDAC6 genes is represented in histograms after oligo interference directed to TDP-43, either individually (siRNA A, B, C or D) or combined (siRNA B+C), normalized to the expression of total α-tubulin. When indicated, *P < 0.01 and **P < 0.05 are P values for Student’s *t* test.

The silencing of TDP-43 decreases the level of the HDAC6 enzyme with subsequent stabilization of the levels of acetylated *α*-tubulin (**Figure 4A**, *HDAC6 and acetylated α-tubulin associated western blot bands*, and **Figure 4B**, *quantified in histograms*), thereby promoting an appropriated cellular environment for HIV-1 viral infection.

We decided to use a combination of two oligos, siRNA-TDP-43 B and C, in order to silence TDP-43. Under this experimental condition, we assayed HIV-1 Env-mediated pore fusion formation and early entry an infection.

We first analyzed how siRNA-TDP-43 silencing affect the ability to the viral Env to promote pore fusion formation, by using the HIV-1 Env-mediated cell-to-cell fusion model. TDP-43 silenced Hela P5 cells were co-cultured with either HeLa 243 (X4-tropic Env) or HeLa ADA (R5-tropic Env) cells, observing an increase of the Env-mediated cell-to-cell fusion activity when compared to scrambled, control cells (**Figure 5**). However, this positive effect on HIV-1 Env-mediated cell-to-cell fusion correlates with low levels of the antifusogenic enzyme HDAC6 in cells were TDP-43 has been knocked-down. The enhancement of the fusogenic activity of HIV-1 Env after TDP-43 knockdown was not related of the viral Env tropism (**Figure 5**).

**Figure 5.**
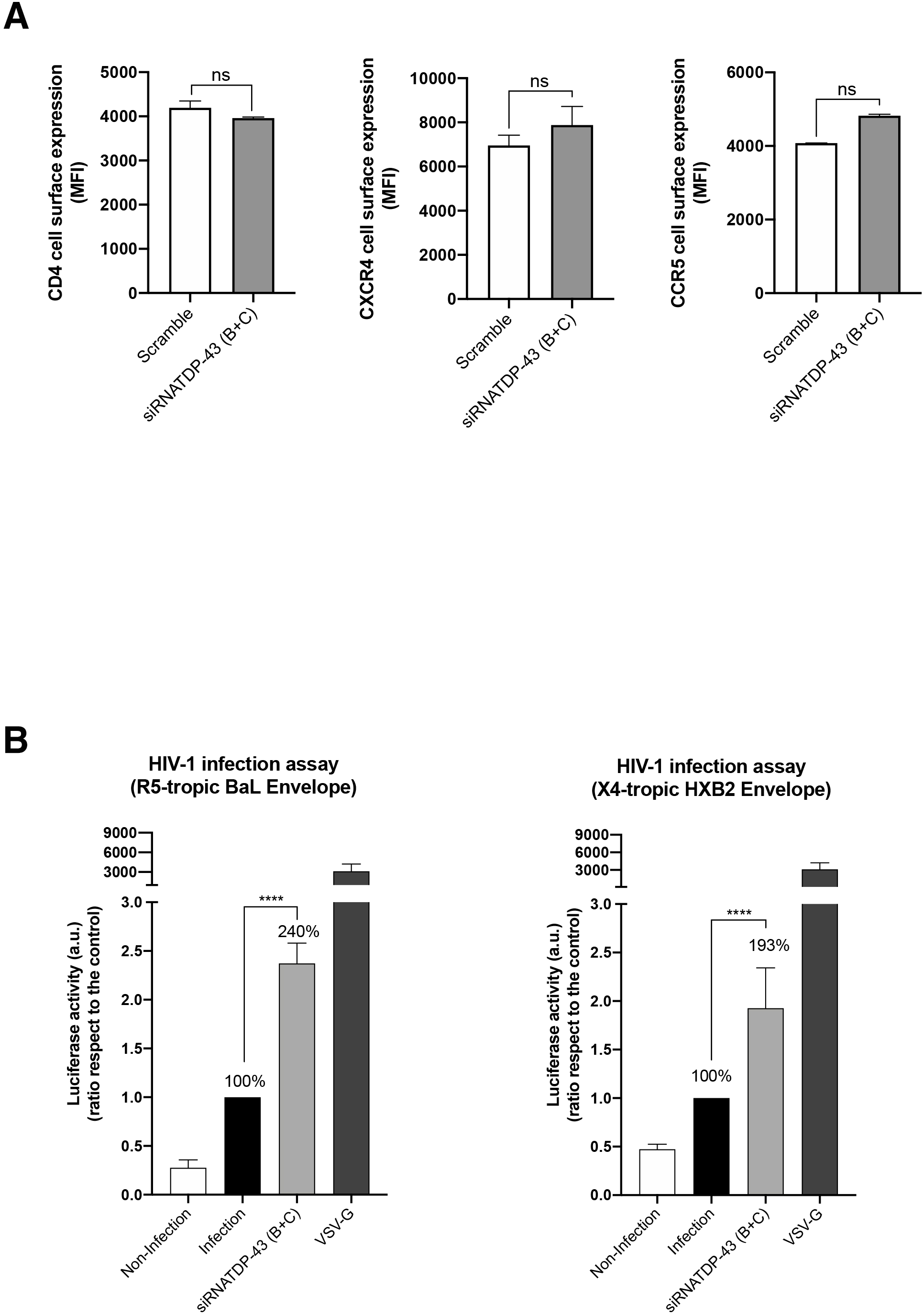
Analysis of CD4 receptor, CXCR4 and CCR5 co-receptors on the cell membrane surface and virus infection with reference envelopes after mRNA silencing directed to endogenous TDP-43. **(A)** Effect of siRNA-TDP-43 oligos on cell-surface expression of HIV-1 receptor and co-receptors (CD4 and CCR5 and CXCR4, respectively) in permissive CEM.NK-CCR5+/CD4+ T cells. Data are mean ± S.E.M. of four independent experiments carried out in triplicate * p < 0.05, ** p < 0.01, *** p < 0.001, Student’s t-test. **(B)** Effects of the four siRNA-TDP-43 oligos in HIV-1 infection with pseudoviruses bearing reference Envs. Viral particles produced with R5-tropic pNL4-3.Luc.R-E-/Env-BaL or /X4-tropic Env-HBX2 viral inputs with CEM.NK.CCR5 permissive T cells previously nucleofected with siRNA-TDP-43 oligos, and assayed in luciferase-quantified viral infection experiments. Data are mean ± S.E.M. of six independent experiments. VSV-G pseudotyped viruses were used as control of viral production and infection.

Next, we infect TDP-43-silenced cells with synchronous doses of HIV-1 (luciferase-reporter pseudovirus) bearing the R5-tropic BaL Env or the X4-tropic HXB2 Env. We observed that specific siRNA silencing of endogenous TDP-43 did not negatively affect the expression of CD4, CCR5 and CXCR4 (**Figure 6**). CEM.NKR-CCR5 T cells were easier to infect when TDP-43 was silenced, expressing low levels of HDAC6 and increased levels of acetylated *α*-tubulin, when compared to control cells (**Figure 6B**). Once again, the effect observed on HIV-1 infection was independent of the viral tropism (**Figure 6B**).

**Figure 6.**
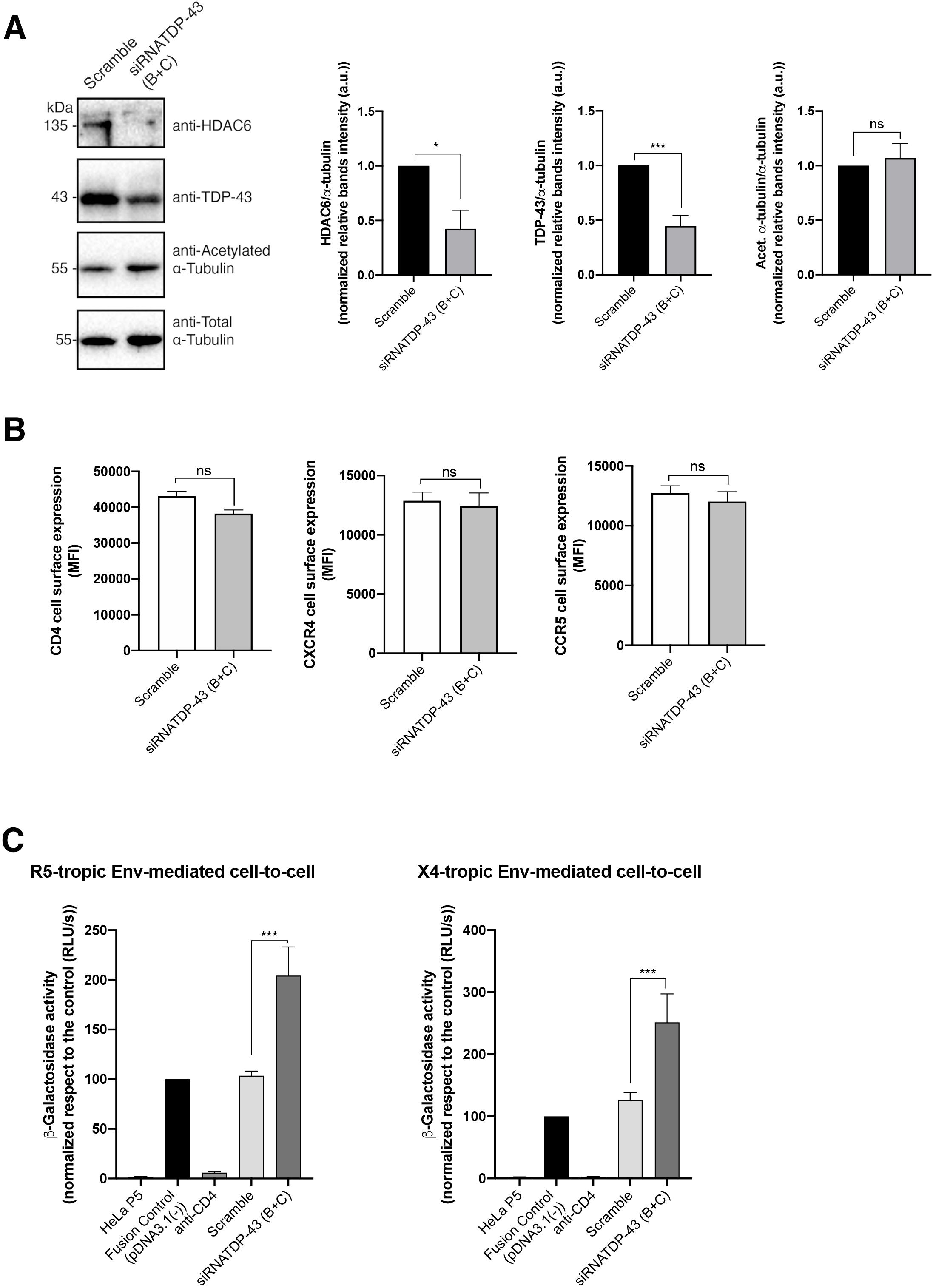
Membrane fusion is favored by the mRNA interference of TDP-43. **(A)** Quantitative western blot analysis of HeLa P5 cell lysate after silencing TDP-43 with specific siRNA oligos, using 1 μM and its effect over endogenous proteins (HDAC6 and acetylated *α*-tubulin). HDAC6/*α*-tubulin and acetylated *α*-tubulin/*α*-tubulin intensity band ratios are shown. **(B)** Effect of silencing of mRNA-TDP-43 in HeLa P5 according to cell-surface expression of viral receptors was analyzed under experimental condition. Data are mean ± S.E.M. of four independent experiments carried out in triplicate * p < 0.05, ** p < 0.01, *** p < 0.001, Student’s t-test. **(C)** Effect of silencing of endogenous TDP-43 in HeLa P5 cells on HIV-1 Env-mediated membrane fusion measured by β-Galactosidase production, and referred to the pcDNA3.1(-) control condition (100% of membrane fusion). Untreated HeLa P5 cells were used as non-fused cells control. Treated HeLa P5 cells were fused with HeLa cells expressing X4-tropic (243) or R5-tropic (ADA) Env, in the presence or absence of a neutralizing anti-fusogenic anti-CD4 mAb.

Therefore, the degree of permissibility to HIV infection of this CD4+ T cells appears to be dependent of the level of TDP-43 which condition the expression level of the antiviral HDAC6 enzyme and MTs decetylation in *α*-tubulin.

### TDP-43 affects the viral function of primary viral Envs from virus of HIV+ individuals with different clinical phenotypes

We would like to seek for the effect of TDP-43 on cell permissibility against primary Env from HIV+ individuals with different clinical phenotypes, such as long-term non-progressors, elite controllers (LTNP-ECs), viremic non-progressors (VNPs) and rapid progressors (RPs) that we have characterized before in their viral function^20, 21^. We observed that HIV-1 Envs from virus of LTNP-EC individuals that control viral infection present less infection activity compared to viral Envs of viremic HIV+ individuals (VNPs) and to those that do not control viral infection at all (RPs), as reported^20, 21^. In Envs from all clinical phenotypes, over-expression of wt-TDP-43 impaired HIV-1 Env-mediated infection (**Figure 7**), revealing that VNP and RP Envs lost their functionality even achieving the levels observed with the inefficient LTNP-EC Envs. On the contrary, specific siRNA-TDP-43 (B+C oligos) knock-down enhanced the infection activity of the HIV-1 Env of VNP and RP individuals (**Figure 8**). Remarkably, TDP-43 silencing renders cells more permissive even to inefficient LTNP-EC Envs (**Figure 8**).

**Figure 7.**
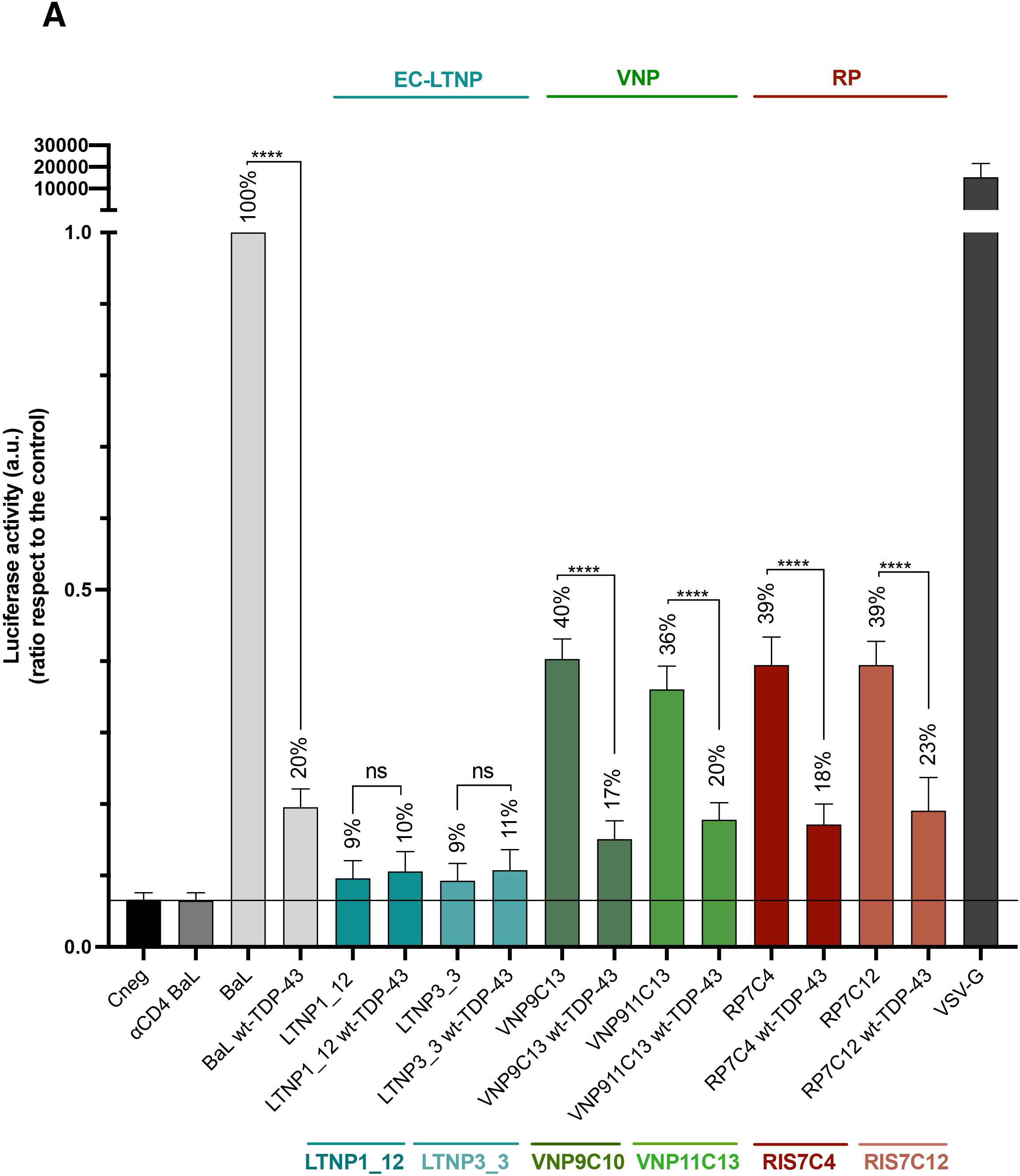
Overexpression of TDP-43 wt construct impairs HIV-1 infection with envelopes from primary viruses isolated from extreme clinical phenotype patients. **(A)** Effect of the overexpression of Flag-wt-TDP-43 construct on the HIV-1 infection with pseudoviruses with R5-tropic pNL4-3.Luc.R-E- and primary viral Envs from virus of LTNP-EC, VNP and RP patients into CEM.NK.CCR5 permissive T cells, previously nucleofected with the TDP-43 construct, and assayed in luciferase-quantified viral infection experiments. Data are mean ± S.E.M. of six independent experiments. VSV-G pseudotyped viruses were used as control of viral production and infection.

**Figure 8.**
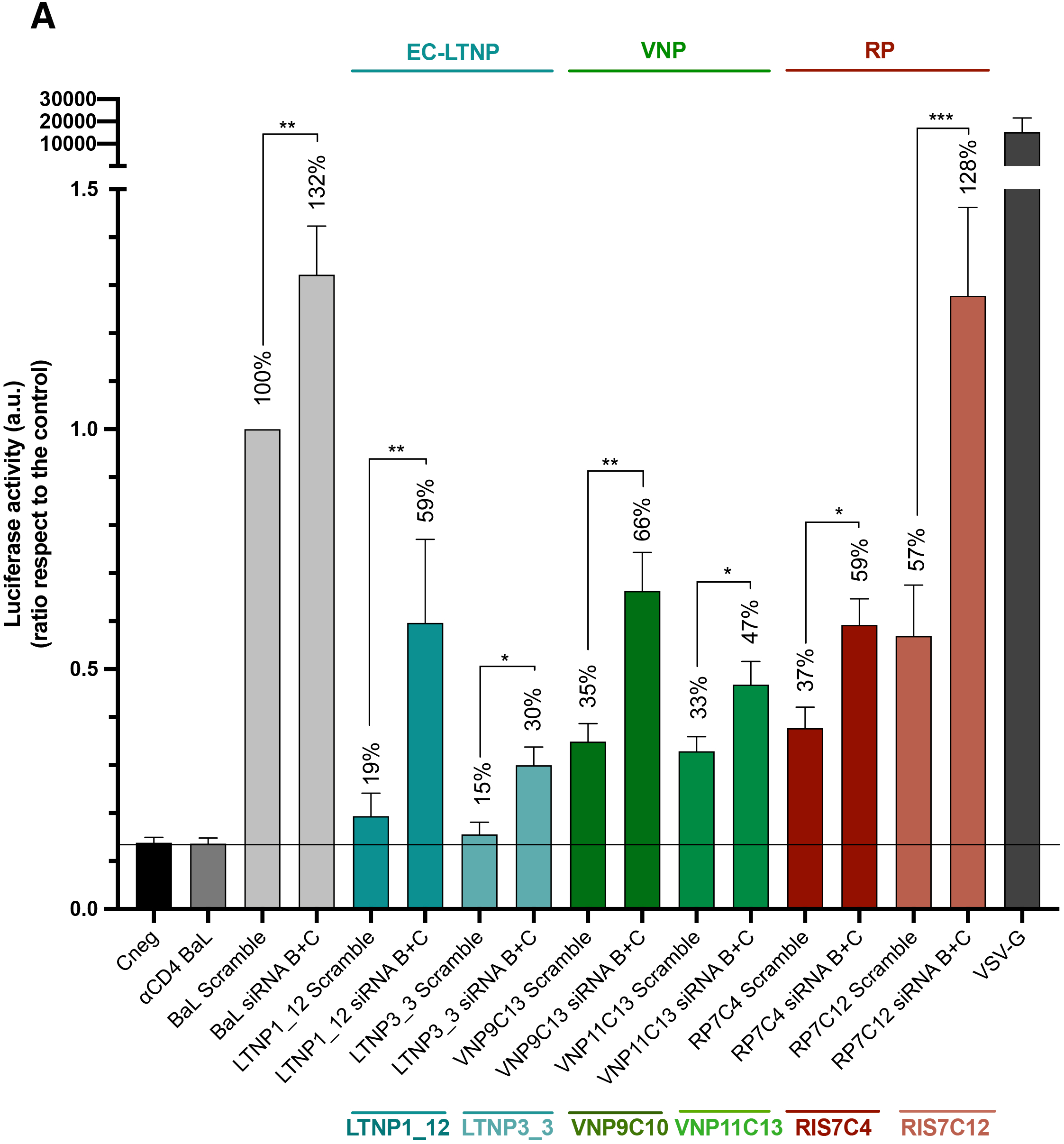
Silencing of endogenous TDP-43 allows and favors HIV-1 infection with envelopes from primary viruses isolated from extreme clinical phenotype patients. **(A)** Effect of the silencing of endogenous TDP-43 on the HIV-1 infection with pseudoviruses with R5-tropic pNL4-3.Luc.R-E- and envelopes from LTNP-EC, VNP and RP patients into CEM.NK.CCR5 permissive T cells previously nucleofected with siRNA-TDP-43 oligos, and assayed in luciferase-quantified viral infection experiments. Data are mean ± S.E.M. of six independent experiments. VSV-G pseudotyped viruses were used as control of viral production and infection.

Altogether these results prompted to suggest that TDP-43 by conditioning HDAC6 mRNA and subsequent protein levels alters the permissive status of target cells against HIV-1 fusion and infection.

## Discussion

In this work, we have observed that over-expression of wt-TDP-43, a cell factor that mainly localizes at the nucleus, stabilized the antiviral enzyme HDAC6, increasing its mRNA and protein levels. Under these experimental conditions, target cells were not efficiently infected by HIV-1 compared to untreated, control cells. This TDP-43- mediated antiviral effect occurred with HIV-1 virus bearing a R5-tropic viral Env as well as an X4-tropic Env. The enhanced levels of the antiviral HDAC6 enzyme in cells over-expressing TDP-43 could be related to the anti-HIV-1 effect exerted by TDP-43. To confirm this TDP-43/HDAC6 antiviral relationship, we further analyzed the ability of HIV-1 Env to promote pore fusion formation in a well characterized cell-to-cell fusion assay using target cells over-expressing wt-TDP-43. These treated cells increased the protein levels of the anti-fusogenic HDAC6 enzyme and when co-cultured with effector cells, stably expressing HIV-1 viral Env at cell-surface, cell-to-cell fusion occurred to a greater extent compared to the same experiment performed with control target cells. It has been reported that level of expression of the tubulin-deacetylase HDAC6 in target cells conditions HIV-1 Env-mediated pore fusion formation and early infection activities^15, 20^. Thus, it is conceivable that TDP-43 negatively regulates this viral Env function by stabilizing HDAC6, an independently of the viral Env tropism. The specific siRNA silencing of endogenous TDP-43 in target cells consistently leads to an increase in the fusogenic and infection activities of the HIV-1 Env. In target cells where endogenous TDP-43 was silenced by using specific siRNA oligos, a significant decrease in the levels of HDAC6 occurred, which favors HIV-1 Env-mediated fusion and infection. Therefore, these results indicate that low levels of TDP-43 enhance target cell permissibility to HIV infection at least by negatively altering the cellular levels of the antiviral HDAC6 enzyme.

These observations were confirmed by using primary Envs from virus of HIV+ individuals with different clinical phenotypes. An increase in the level of expression of wt-TDP-43 strongly diminished the infection activity of HIV-1 Envs from virus of VNP and RP HIV+ individuals, even reaching the levels of the inefficient HIV-1 Env of virus of LTNP-EC individuals. On the contrary, low levels of endogenous TDP-43, obtained after siRNA knocking-down, significantly favors the infection activity of primary HIV-1 Envs of VNP and progressors individuals, even observing an increase in the infection ability of the primary HIV-1/LTNP-EC Envs. This fact clearly points out that TDP-43 regulates cell permissibility to HIV infection by affecting vial Env fusion and infection capacities, thereby altering the cellular level of the HDAC6 antiviral factor. In this regard, it has been reported that the ability of the viral Env to trigger signals that overcome the HDAC6 barrier is directly related to its fusion and infection activities^15, 20^.

Thus, TDP-43 seems to regulate cell permissibility to HIV infection. This event may have negative consequences in HIV+ LTNP-EC individuals, particularly if a negative regulation of TDP-43 may occur with a concomitant decrease in HDAC6 that render cells more permissive against inefficient LTNP-EC Envs thereby favoring viral infection. This could be another factor that worth to be explored, apart from the immune responses, in EC individuals that loss their status of natural controllers of the infection^51–53^. In the central nervous system (CNS), the unbalance between TDP-43 and HDAC6 levels of expression and functions seems to be related with neurodegeneration and disease^5–12^. In this matter, it has been reported that HDAC6 prevents the neurotoxicity of the HIV-1-Env gp120 subunit in cortical neurons^54^. Maybe the TDP-43/HDAC6 axis in the CNS could act protecting against HIV-1 toxicity in this tissue, whereas an unbalance in their expression levels and functions could favor HIV-1 damage, but this needs to be explored further.

Therefore, TDP-43 appears to regulate cell permissibility to HIV-1 infection by conditioning viral Env-mediated fusion and infection capacities by modulating the level of expression of the antiviral enzyme HDAC6.

## Author contributions

R-CR wrote the paper and performed the research and supervised experimental design; S-PY, R-GM, JM-LS, J-EH, J-GL, A-IC, LA-RR and A-MB performed the research and supervised experimental design; R-TG, R-DG, CC and MP performed statistics and the research; JB, C-F and JM-LS designed the research and laboratory experiments, supervised experimental design, analysis and interpretation of data. A-VF wrote the paper, conceived and designed the research and laboratory experiments, supervised experimental design, analysis and interpretation of data. All authors read and approved the final manuscript.

